# β-blockers augment L-type Ca^2+^ channel activity by target-selective spatially restricted β_2_AR-cAMP-PKA signaling in neurons

**DOI:** 10.1101/668913

**Authors:** Ao Shen, Dana Chen, Manpreet Kaur, Bing Xu, Qian Shi, Joseph M. Martinez, Kwun-nok Mimi Man, Johannes W. Hell, Manuel F. Navedo, Xi-Yong Yu, Yang K. Xiang

## Abstract

G protein-coupled receptors (GPCRs) transduce pleiotropic intracellular signals in mammalian cells. Here, we report that some antagonists of β adrenergic receptors (βARs) such as β-blocker carvedilol and alprenolol activate β_2_AR at nanomolar concentrations, which promote G protein signaling and cAMP/PKA activity without action of G protein receptor kinases (GRKs). The cAMP/PKA signal is restricted within the local plasma membrane domain, leading to selectively enhance PKA-dependent augment of endogenous L-type calcium channel (LTCC) activity but not AMPA receptor in hippocampal neurons. Moreover, we have engineered a mutant β_2_AR that lacks serine 204 and 207 in the catecholamine binding pocket. This mutant can be preferentially activated by carvedilol but not the orthosteric agonist isoproterenol. Carvedilol activates the mutant β_2_AR in hippocampal neurons augmenting LTCC activity through cAMP/PKA signaling. Together, our study identifies a mechanism by which β-blocker-dependent activation of GPCRs at low ligand concentrations promotes local cAMP/PKA signaling to selectively target membrane downstream effectors such as LTCC in neurons.

## Introduction

GPCRs often signal not only through canonical G proteins but also through noncanonical G protein-independent signaling, frequently via G protein receptor kinases (GRKs) and β-arrestins (1, 2). One of the universal features of GPCRs is that they undergo ligand-induced phosphorylation at different sites by either GRKs or second messenger dependent protein kinases such as protein kinase A (PKA). The phosphorylated GPCRs thus may present distinct structural features that favor receptor binding to different signaling partners, engaging distinct downstream signaling cascades (3-5). Some ligands can differentially activate a GPCR via a phenomenon known as functional selectivity or biased signaling (6, 7). For example, stimulation of β_2_-adrenergic receptor (β_2_AR), a prototypical GPCR involved in memory and learning in the central nervous system (CNS) and regulation of metabolism and cardiovascular function, promotes phosphorylation by both GRKs and PKA (8-11). We have recently identified spatially segregated subpopulations of β_2_AR undergoing exclusive phosphorylation by GRKs or PKA in a single cell, respectively. These findings indicate specific GPCR subpopulation-based signaling branches can co-exist in a single cell (12). GRK-mediated phosphorylation promotes pro-survival and cell growth signaling via β-arrestin-dependent mitogen-activated protein kinase (MAPK/ERK) pathways, prompting the search for biased ligands that preferentially activate β-arrestin pathways (13-18). On the other hand, our recent studies show that the cAMP/PKA-dependent phosphorylation of β_2_AR controls ion channel activity at the plasma membrane in primary hippocampal neurons (12).

β-blockers are thought to reduce cAMP signaling because they either reduce basal activity of βARs or block agonist-induced receptor activation. While β-blockers are successful in clinical therapies of a broad range of diseases, their utility is limited by side effects in both the CNS and peripheral tissues (19, 20). Indeed, studies have revealed that some β-blockers display partial agonism and can promote receptor-Gs coupling at high concentrations *in vitro* (21-23). Accordingly, some β-blockers display intrinsic properties mimicking sympathetic activation (sympathomimetic β-blockers) (24-26). The mechanism remains poorly understood because classic cAMP assay do not show even minimal cAMP signal induced by these β-blockers (24, 25).

In this study, we show that the β-blockers carvedilol and alprenolol can promote Gs protein coupling to β_2_AR and cAMP/PKA but not GRK activity at nanomolar concentrations. Thus these β-blockers are emerging as partial agonists rather than strict antagonists in mammalian cells. This cAMP/PKA signaling is spatially restricted, selectively promoting phosphorylation of β_2_AR and Ca_V_1.2 by PKA which augments LTCC activity in primary hippocampal neurons. Furthermore, we have engineered a mutant β_2_AR that can be selectively activated by carvedilol but not by the orthosteric agonist isoproterenol (ISO) to stimulate PKA but not GRK. Together, these studies identify a unique mechanism by which β-blockers activate β_2_AR at low concentrations, which promotes Gs/cAMP/PKA signaling branch and selectively targets downstream LTCC channels in neurons. This observation may also explain sympathomimetic effects of β-blockers in the CNS.

## Results

### AR-mediated PKA-phosphorylation of β_2_AR in HEK293 cells

In this study, we applied two sets of well-characterized phospho-specific antibodies, anti-pS261/262 and anti-pS355/356 to examine a series of β-blockers for their effects on the phosphorylation of β_2_AR at its PKA and GRKs sites, respectively (12, 27, 28). We found that various β-blockers including alprenolol (ALP), carvedilol (CAR), propranolol (PRO) and CGP12177 (177) were able to stimulate phosphorylation of β_2_AR at PKA sites expressed in HEK293 cells, similar to the βAR agonist isoproterenol (ISO) (**Fig. 1a**). In contrast, other β-blockers, i.e., ICI118551 (ICI), timolol (TIM) and metoprolol (MET), were not able to do so (**Fig. 1a**). The ligand-induced phosphorylation of β_2_AR was blocked by β_2_AR-specific antagonist ICI but not β_1_AR-specific antagonist CGP20712A (CGP) (**Fig. 1b,c**). We chose CAR and ALP for further study. We found that CAR and ALP promoted phosphorylation of β_2_AR by PKA even at nanomolar concentrations (**Fig. 2a,b**), which was paralleled by concentration-dependent increases in phosphorylation of ERK (**Supplemental Fig. 1**). The roles of β_2_AR and PKA in this phenomenon were confirmed by inhibition of β_2_AR with ICI and inhibition of PKA with H89, respectively (**Fig. 2c,d**). In contrast, those β-blockers induced at best minimal increases in phosphorylation of β_2_AR at GRK sites and only at high concentrations, consistent with previous report (29) (**Fig. 1a** and **Supplemental Fig. 1,2**). As positive control, the βAR agonist ISO promoted robust increases in both PKA and GRK phosphorylation of the receptors at different concentrations ranging from nanomolar to micromolar (**Fig. 1,2** and **Supplemental Fig. 1,2**). In the CNS, β_2_AR emerges as a prevalent postsynaptic norepinephrine effector at glutmatergic synapses (30-33). Consistent with the data from HEK293 cells, we found that β-blockers CAR and ALP activated β_2_AR and promoted phosphorylation of the receptor by PKA in hippocampal neurons (**Fig. 4a**). Together, these data suggest that certain β-blockers selectively promote PKA phosphorylation of β_2_AR in HEK293 and primary hippocampal neurons.

**Figure 1.**
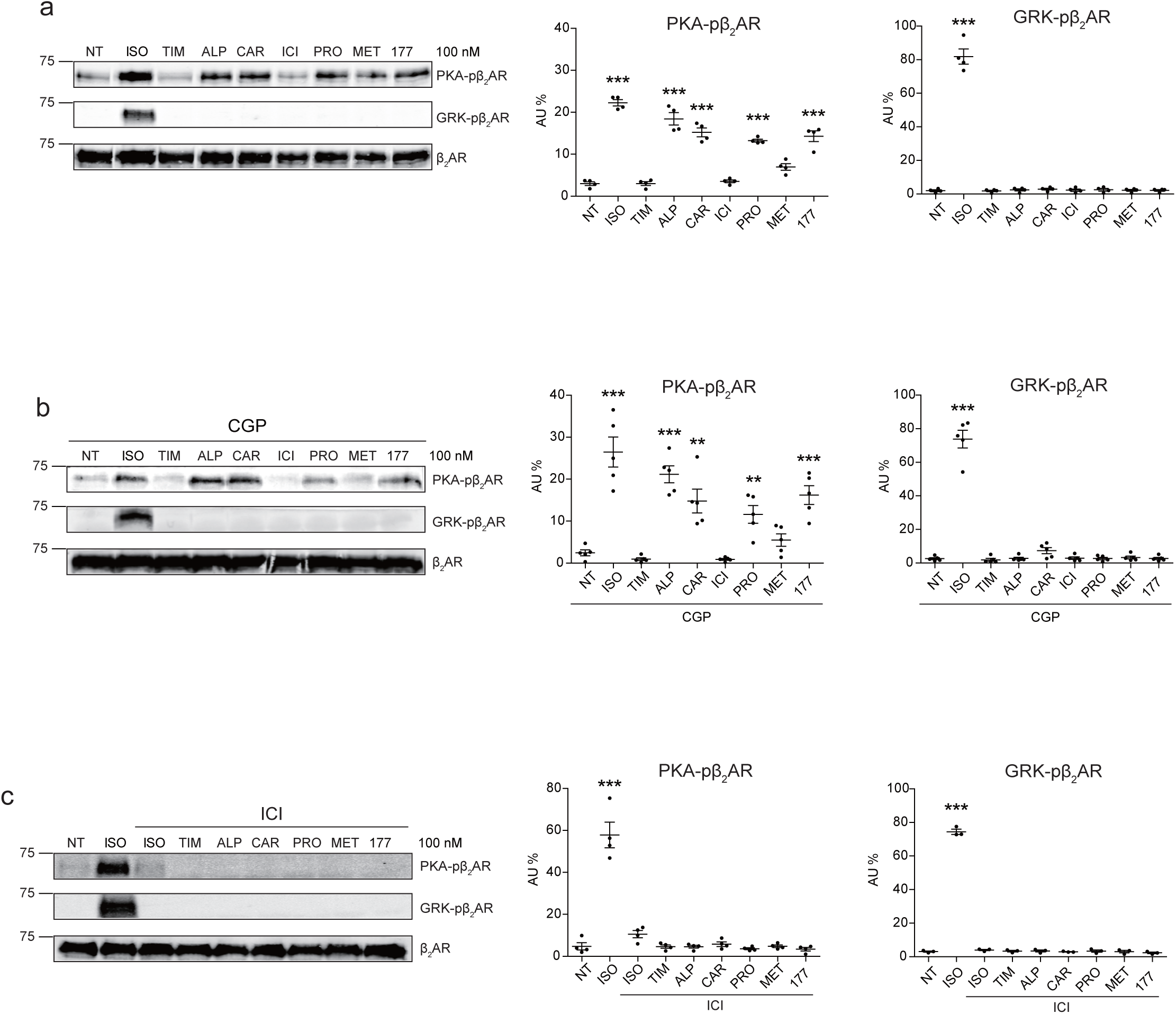
Carvedilol and alprenolol selectively promote phosphorylation of β_2_AR at PKA sites. HEK293 cells stably expressing FLAG-tagged β_2_AR were either directly stimulated for 5 minutes with the βAR agonist ISO and different β-blockers at indicated concentrations (**a**), or pretreated for 15 minutes with 1 μM β_1_AR antagonist CGP20712A (**b**) or 10 μM β_2_AR antagonist ICI118551 (**c**) before the treatment. The phosphorylation levels of β_2_AR on its PKA and GRK sites were determined by Western blot with phosphor-specific antibodies, and signals were normalized to total β_2_AR detected with anti-FLAG antibody. NT, no treatment; ISO, isoproterenol; TIM, timolol; ALP, alprenolol; CAR, carvedilol; ICI, ICI118551; PRO, propranolol; MET, metoprolol; 177, CGP12177; CGP, CGP20712A. Error bars denote s.e.m., P values are computed by one-way ANOVA followed by Tukey’s test between NT and other groups.

**Figure 2.**
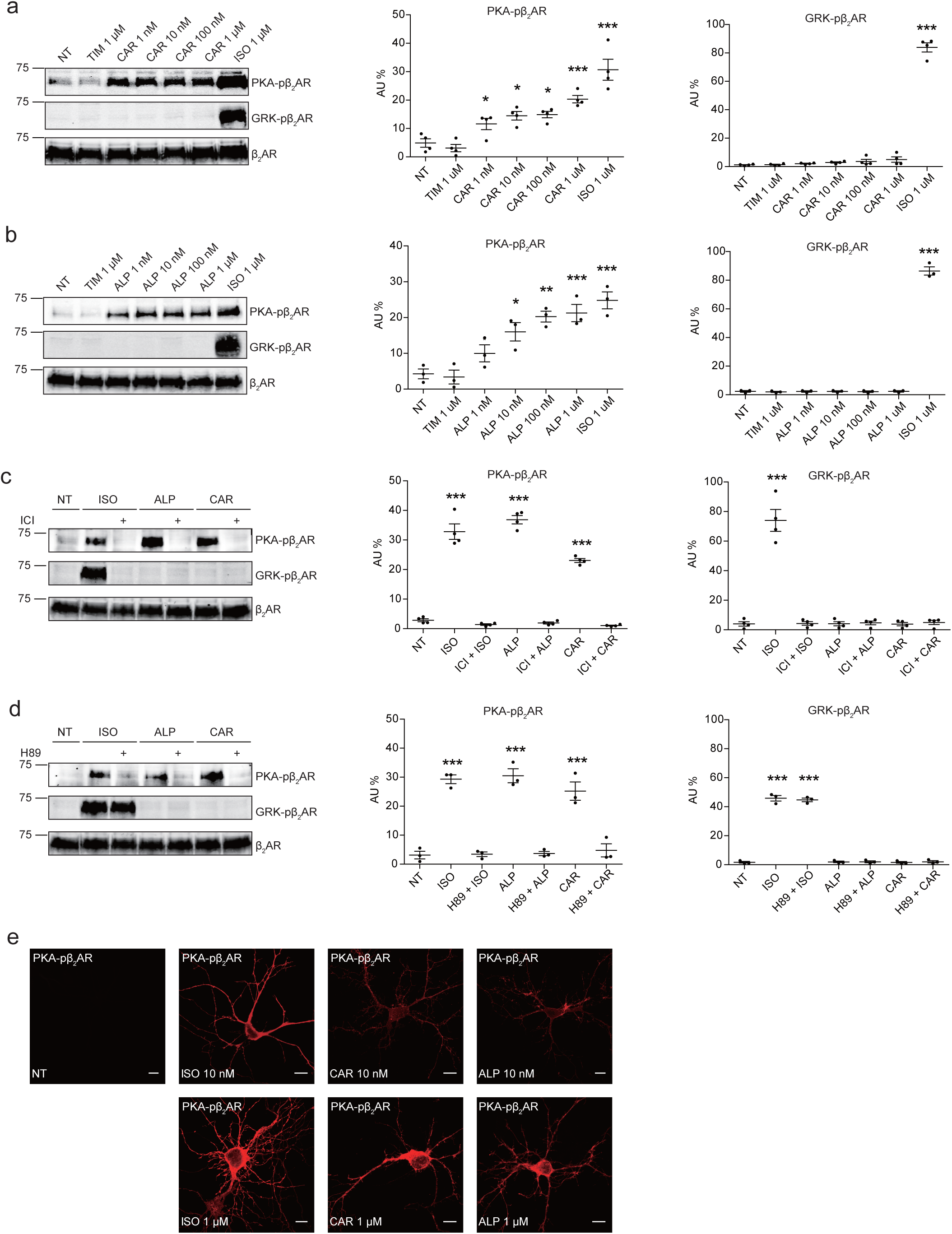
Carvedilol and alprenolol induce concentration-dependent PKA phosphorylation of β_2_AR in HEK293 and hippocampal neurons. HEK293 cells stably expressing FLAG-tagged β_2_AR were treated with increasing concentrations of CAR (**a**) and ALP (**b**) or pretreated for 15 minutes with 10 μM β_2_AR antagonist ICI118551 (**c**) and PKA inhibitor H89 (**d**) before stimulated with 1 μM indicated drugs for 5 minutes. The phosphorylation levels of β_2_AR on its PKA and GRK sites were determined with phosphor-specific antibodies, and signals were normalized to total β_2_AR detected with anti-FLAG antibody. Experiments were performed in the presence of 1 μM β_1_AR-selective antagonist CGP20712A to block endogenous β_1_AR signaling. NT, no treatment; ISO, isoproterenol; ALP, alprenolol; CAR, carvedilol; ICI, ICI118551. Error bars denote s.e.m., P values are computed by one-way ANOVA followed by Tukey’s test between NT and other groups. **e** Rat hippocampal neurons expressing β_2_AR were treated for 5 minutes with 10 nM or 1 μM indicated drugs on 12 DIV and immuno-stained for PKA-phosphorylated β_2_AR. Confocal images show PKA-phosphorylated β_2_AR in agonist- or β-blocker-stimulated neurons have similar distribution. Scale bar, 10 μm. Representative of 6 images for each condition, three experiments.

**Figure 3.**
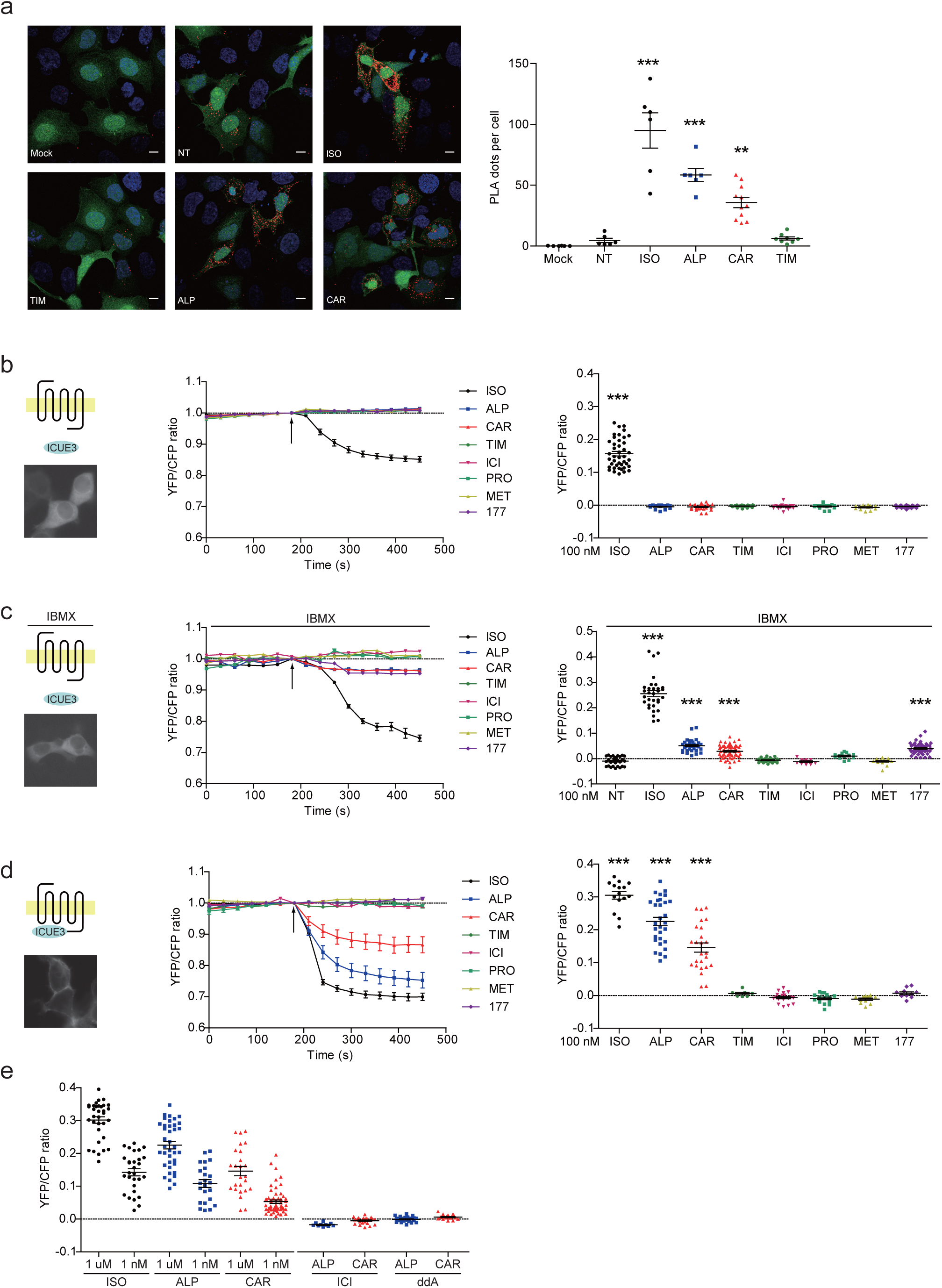
Carvedilol and alprenolol promote Gsα recruitment to β_2_AR and increase locally restricted cAMP signal. **a** HEK293 cells co-expressing FLAG-tagged β_2_AR, HA-tagged Gsα and EGFP were stimulated with 100 nM ISO or indicated β-blockers for 5 minutes. In proximity ligation assay (PLA), cells were immuno-stained with HA and β_2_AR antibody, nuclei were counterstained with DAPI. The green EGFP signal represents transfected cells, and red PLA signal represents Gsα and β_2_AR interactions. Carvedilol and alprenolol promoted Gsα recruitment to β_2_AR, but timolol could not. Scale bar, 10 μm. Representative of n= 6, 6, 6, 6, 11 and 8 images respectively, three experiments. **b**-**c** HEK293 cells expressing ICUE3 biosensor were treated with 1 μM ISO or indicated β-blockers (**b**), or together with 100 μM phosphodiesterase inhibitor IBMX (**c**). **d**-**e** HEK293 cells expressing β_2_AR-ICUE3 biosensor were treated with indicated concentration of ISO or β-blockers. In some cases, cells were pretreated for 30 minutes with the β_2_AR antagonist ICI (10 μM) or the adenylate cyclase inhibitor ddA (50 μM) before adding β-blockers. Changes in ICUE3 FRET ratio (an indication of cAMP activity) were measured. Experiments were performed in the presence of 1 μM β_1_AR-selective antagonist CGP20712A to block endogenous β_1_AR signaling. Mock, no primary antibody; NT, no treatment; ISO, isoproterenol; TIM, timolol; ALP, alprenolol; CAR, carvedilol; ICI, ICI118551; PRO, propranolol; MET, metoprolol; 177, CGP12177, IBMX, 3-isobutyl-1-methylxanthine; ddA, 2’,5’-dideoxyadenosine. Error bars denote s.e.m., P values are computed by one-way ANOVA followed by Tukey’s test between NT (**a**) or TIM (**b**-**e**) and other groups.

**Figure 4.**
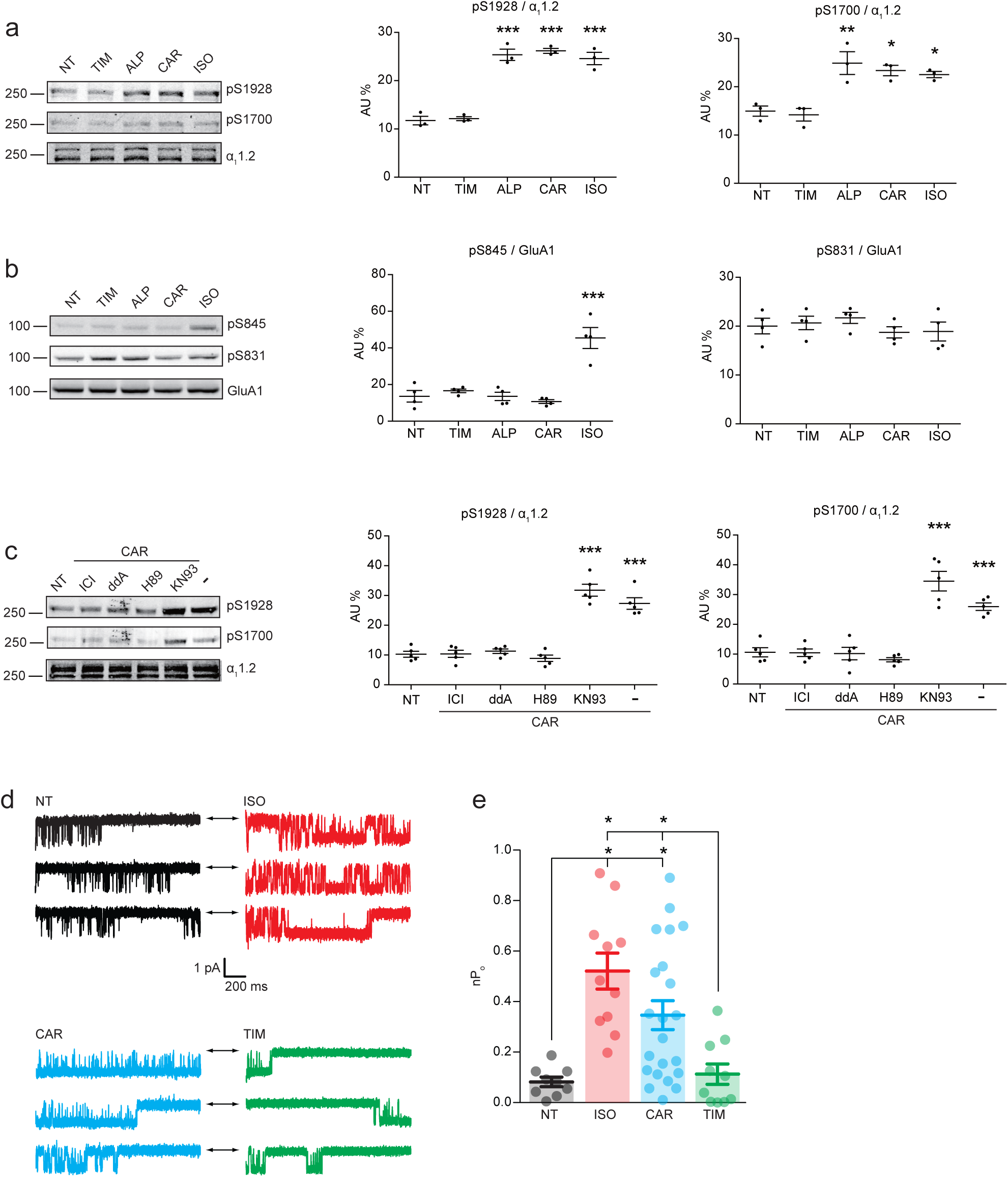
Carvedilol promotes endogenous β_2_AR-dependent phosphorylation of LTCC α_1_1.2 by PKA and augment of channel activity in hippocampal neurons. **a** Rat neurons on 10-14 DIV were treated for 5 minutes with 1 μM indicated drugs. The phosphorylation of endogenous LTCC α_1_1.2 subunit was determined with phospho-specific antibodies and normalized to total α_1_1.2. **b** Neurons on 10-14 DIV were treated for 5 minutes with 1 μM indicated drugs. The phosphorylation of endogenous AMPAR GluA1 subunit was determined with phosphor-specific antibodies, and signals were normalized to total GluA1.**c** Neurons were pretreated for 30 minutes with 10 μM β_2_AR inhibitor ICI, 50 μM AC inhibitor ddA, 10 μM PKA inhibitor H89 or 10 μM CaMKII inhibitor KN93 and then stimulated with 1 μM CAR for 5 minutes. Carvedilol-induced LTCC phosphorylation depends on endogenous β_2_AR, AC and PKA, but not CaMKII. **d** Representative single-channel recordings of endogenous LTCC Ca_V_1.2 currents in rat hippocampal neurons on 7-10 DIV after depolarization from −80 to 0mV without (NT) and with stimulation of 1 μM indicated drugs in the patch pipette. Arrows throughout the figure indicate the 0-current level (closed channel). **e** The overall channel activity (nPo) of Ca_V_1.2 was quantified from **d**. Error bars denote s.e.m., P values are computed by one-way ANOVA followed by Tukey’s test between NT and other groups in **a**-**c** and Mann-Whitney test in **e**.

### Carvedilol and alprenolol promote Gsα recruitment to β_2_AR and increase locally restricted cAMP signal

The western blot data on PKA phosphorylation of β_2_AR indicates a stimulation of the receptor-mediated Gs/AC/cAMP pathway by these β-blockers. We measured ligand-induced Gsα recruitment to β_2_AR with an *in situ* proximity ligation assay (PLA), which allows direct visualization and quantification of protein-protein interactions. We showed that ISO, CAR and ALP were able to increase the PLA signals between β_2_AR and Gsα, indicating recruitment of Gsα to β_2_AR (**Fig. 3a**). As control, TIM had no effect on the recruitment of Gsα to β_2_ARs. The role of Gs/AC in CAR-induced PKA phosphorylation of β_2_AR was further validated by AC-specific inhibition with 2’,3’-dideoxyadenosine (ddA, **Supplemental Fig. 3**). These data indicate that CAR and ALP are able to stimulate β_2_AR-Gs signal to increase PKA phosphorylation of the receptor.

β-blockers have been thought to generally block β_2_AR-induced cAMP signal. We hypothesized that the cAMP signal induced by β-blockers is restricted to local plasma membrane domains containing activated receptor, which is not detectable with traditional cAMP assays likely due to limited sensitivity. We applied highly sensitive FRET-based biosensor ICUE3 to detect the dynamics of cAMP signal in living cells (34, 35). The full agonist ISO promoted cAMP signal in HEK293 cells while all β-blockers failed to do so (**Fig. 3b**), in agreement with the classic definition of β-blockers. However, when cells were treated with non-selective phosphodiesterase (PDE) inhibitor IBMX, CAR, ALP and CGP12177 were able to induce small but significant cAMP signal in HEK293 cells (**Fig. 3c**), indicating a role of PDE in suppressing and restricting the distribution of cAMP in the cells. When β_2_AR was exogenously expressed in HEK293 cells, CAR and ALP were able to induce cAMP signal in HEK293 cells even without PDE inhibition (**Supplemental Fig. 4**), probably due to insufficient cAMP-hydrolytic activity of endogenous PDEs to counter cAMP production induced from overexpressed β_2_AR. We then engineered a targeted cAMP biosensor by fusing the biosensor ICUE3 to the C-terminus of β_2_AR (β_2_AR-ICUE3), aiming to detect increases of cAMP within the local domain of activated β_2_AR. CAR and ALP promoted cAMP activity within the immediate vicinity of the receptor. CAR and ALP promoted cAMP signals within the local domain of activated β2AR even at nanomolar concentrations (**Fig. 3d,e**). The local increases of cAMP were abolished by inhibition of β_2_AR with ICI or inhibition of ACs with ddA (**Fig. 3e**). These data confirm that CAR and ALP promote cAMP/PKA activity within the local domain of activated β_2_AR, in contrast to the broad distribution of cAMP/PKA activities induced by ISO in the cells.

### Carvedilol augments the endogenous β_2_AR-dependent PKA phosphorylation of Ca_V_1.2 and its channel activity in hippocampal neurons

The local cAMP signals possess the potential to selectively regulate downstream effectors in receptor complexes or within the vicinity of activated receptors. In the CNS, β_2_AR emerges as a prevalent postsynaptic norepinephrine effector at glutmatergic synapses, where β_2_AR functionally interacts with AMPA receptor (AMPAR) and L-type Ca^2+^ channel (LTCC) Ca_V_1.2, and regulates neuronal excitability and synaptic plasticity (30-33). CAR and ALP, but not TIM significantly increased PKA phosphorylation of S1928 and S1700 of central α_1_1.2 subunit of Ca_V_1.2 in hippocampal neurons when both β_2_AR and LTCC were endogenously expressed (**Fig. 4a**). However, CAR and ALP failed to promote phosphorylation of the AMPAR subunit GluA1 on its PKA site serine 845 (**Fig. 4b**). Like Ca_V_1.2, AMPARs are associated with β_2_AR, Gs, AC and PKA (32-35). These results indicate high selectivity in targeting downstream substrates by this β-blocker-induced signaling in hippocampal neurons. Meanwhile, the CAV and ALP-induced PKA phosphorylation of LTCC were blocked by β_2_AR inhibitor ICI, AC inhibitor ddA, and PKA inhibitor H89, but not CaMKII inhibitor KN93, validating the activation of β_2_AR-cAMP-PKA pathway (**Fig. 4c**). We then examined the effects of CAR on PKA-dependent activation of LTCC Ca_V_1.2 channels using single channel recordings in hippocampal neurons. Consistent with the phosphorylation data, CAR but not TIM significantly increased open probabilities of endogenous Ca_V_1.2 in hippocampal neurons (**Fig. 4d,e**). These data indicate that CAR promotes local cAMP/PKA activities for selective augmentation of LTCC activities in neurons.

### Carvedilol but not isoproterenol selectively activates a mutant β_2_AR to augment LTCC activity in neurons

Structure-functional analyses of β_2_AR have previously revealed distinct residues important for binding to catecholamines and β-blockers (36-39). We hypothesized that mutation of Ser204 and Ser207 sites within β_2_AR binding pocket would abolish receptor hydrogen bonds with the catecholamine phenoxy moieties, thus reducing binding affinity to agonist ISO while having no effect on β-blocker binding (**Fig. 5a**). Such a mutant β_2_AR could thus be selectively activated by CAR. We co-expressed the cAMP biosensor ICUE3 together with either wild-type (WT) β_2_AR or mutant S204A/S207A β_2_AR in MEF cells lacking endogenous β_1_AR and β_2_AR (DKO) to detect receptor signaling induced by different ligands. The mutant S204A/S207A β2AR induced a moderate cAMP signal at high but not low concentrations of ISO (**Fig. 5b**). In contrast, after stimulation with CAR, the β_2_AR mutant S204A/S207A promoted significant cAMP signals at nanomolar concentrations; the overall concentration response curve was similar to those induced by WT β_2_AR (**Fig. 5b**). Accordingly, the ISO-induced PKA phosphorylation of β_2_AR S204A/S207A mutant was selectively abolished at nanomolar concentrations. At higher concentrations, ISO was able to induce reduced PKA phosphorylation of the β_2_AR S204A/S207A mutant when compared to WT β2AR, consistent with the data of cAMP signals (**Fig. 5c**). Meanwhile, ISO failed to induce GRK phosphorylation of β_2_AR S204A/S207A mutant at different concentrations (**Fig. 5c**). In comparison, CAR induced equivalent PKA phosphorylation of β_2_AR WT and S204A/S207A mutant at different concentrations (**Fig. 5d**). These data suggest that CAR, but not ISO selectively activates the S204A/S207A mutant β_2_AR at nanomolar concentrations. We then tested the effects of β_2_AR S204A/S207A mutant on LTCC channel activity after treatment with CAR in hippocampal neurons. In DKO neurons expressing the mutant S204A/S207A β_2_AR, CAR, but not ISO (30 nM) promoted PKA phosphorylation of LTCC α_1_1.2 (**Fig. 6a** and **Supplemental Fig. 5**). In agreement, CAR, but not ISO significantly increased open probabilities of Ca_V_1.2 (**Fig. 6b,c**). Together, CAR but not ISO selectively activates the S204A/S207A mutant β_2_AR at low concentrations and increases channel opening probabilities.

**Figure 5.**
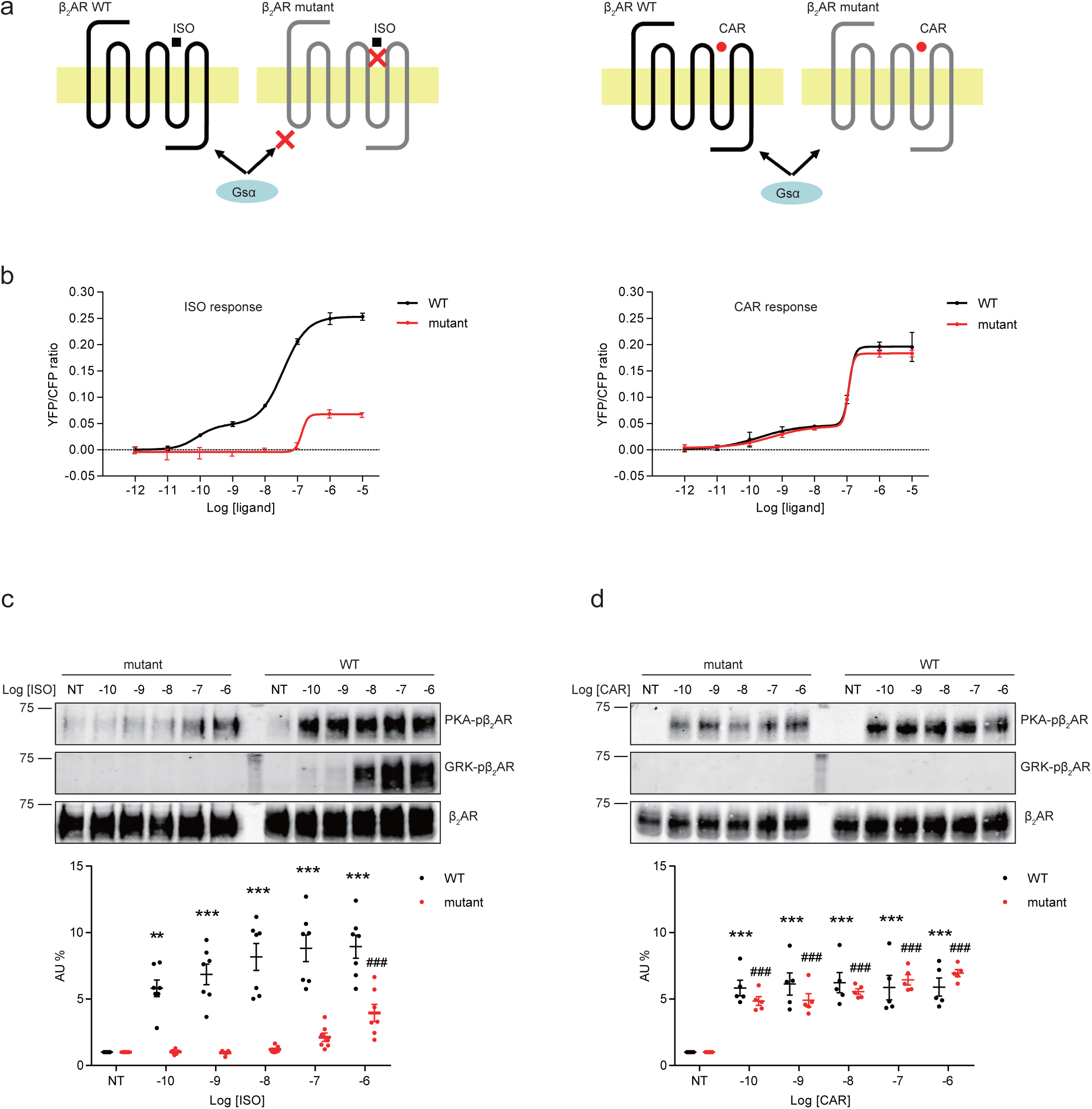
A mutant β_2_AR is selectively activated by carvedilol but not isoproterenol. **a** Schematic of an engineered β_2_AR with S204/207A double serine mutations that loses high affinity binding to ISO but not CAR at nanomolar range. **b** cAMP biosensor ICUE3 and β_2_AR wild-type (WT) or mutant were co-expressed in MEF cells lacking both β_1_AR and β_2_AR. Changes of cAMP FRET ratio by increasing concentrations of ISO or CAR were measured. **c**-**d** HEK293 cells stably expressing FLAG-tagged β_2_AR WT or mutant were stimulated for 5 minutes with increasing concentrations of ISO (**c**) or CAR (**d**). The phosphorylation of β_2_AR on its PKA and GRK sites were determined by Western blot with phospho-specific antibodies, and signals were normalized to total β_2_AR detected with anti-FLAG antibody. Experiments were performed in the presence of 1 μM β_1_AR-selective antagonist CGP20712A to block endogenous β_1_AR signaling. NT, no treatment; ISO, isoproterenol; CAR, carvedilol. Error bars denote s.e.m., P values are computed by one-way ANOVA followed by Tukey’s test between NT and other concentrations.

**Figure 6.**
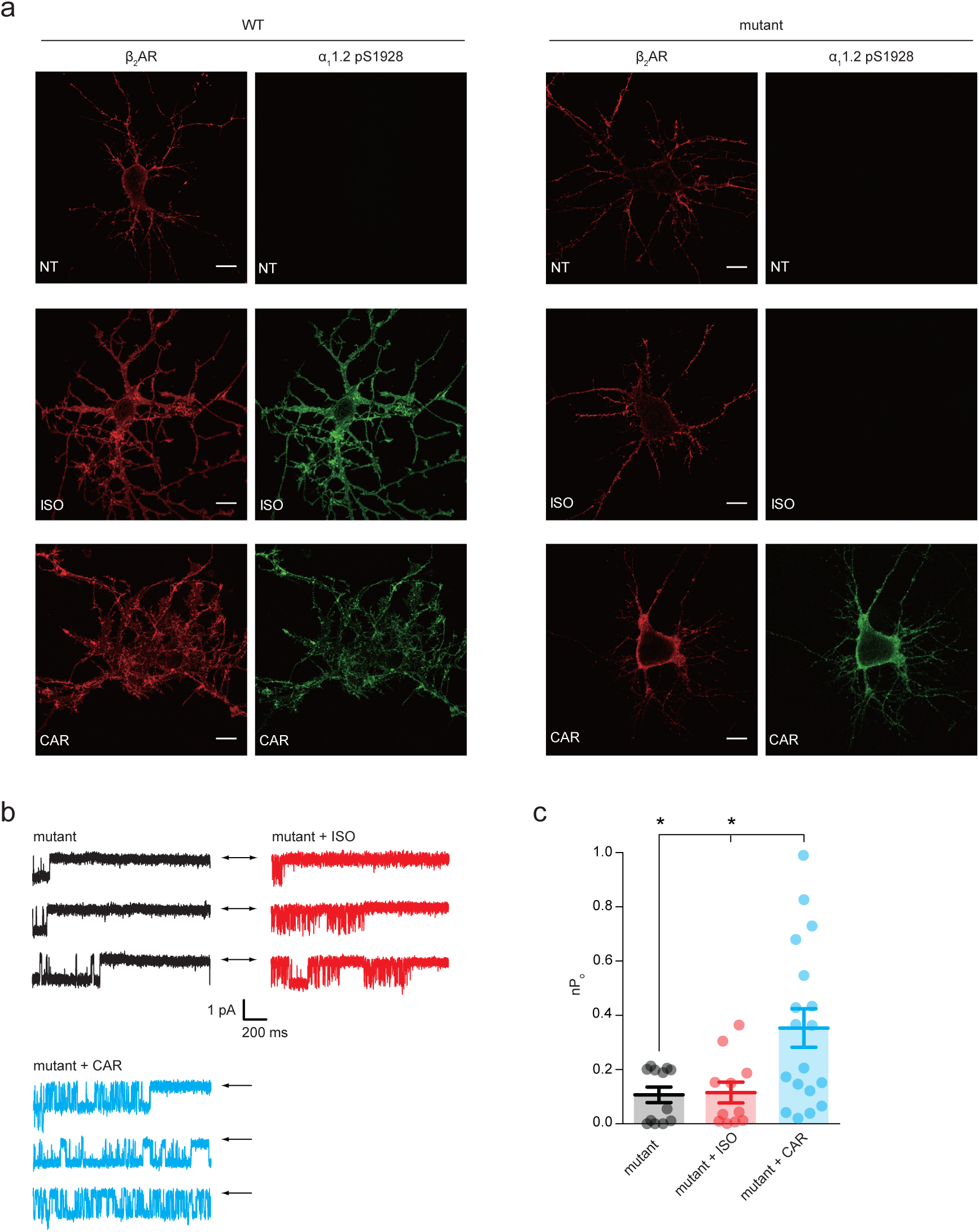
Carvedilol selectively activates the mutant β_2_AR and promotes LTCC activity in hippocampal neurons. **a** β_1_AR/β_2_AR double knockout (DKO) mouse hippocampal neurons on 7-10 days in vitro (DIV) were cotransfected with FLAG-tagged β_2_AR WT or mutant and HA-tagged LTCC α_1_1.2 subunit, 24 hours later cells were either mock treated (upper panel), or treated for 5 minutes with 10 nM ISO (middle panel) or 10 nM CAR (lower panel), fixed and labeled with anti-FLAG and a phosphor-specific antibody for S1928 phosphorylated α_1_1.2. Representative confocal images show mutant β_2_AR lost the ability of promoting LTCC phosphorylation upon ISO stimulation but remained the ability upon CAR stimulation in mouse neurons. Scale bar, 10 μm. Representative of 6 images for each condition, three experiments. **b** Representative single-channel recordings of LTCC Ca_V_1.2 currents in DKO neurons on 7-10 DIV expressing mutant β_2_AR after depolarization from −80 to 0mV without (black traces) and with stimulation of 30 nM indicated drugs in the patch pipette. Arrows throughout the figure indicate the 0-current level (closed channel). **c** The overall channel activity (nPo) of LTCC Ca_V_1.2 was quantified from **b**. Error bars denote s.e.m. with Mann-Whitney test.

## Discussion

In a classic view, agonist stimulation promotes both PKA and GRK phosphorylation of activated GPCRs. In this study using a combination of highly sensitive tools such as engineered FRET-based cAMP sensors and single channel recording together with detection with phospho-specific antibodies, we show that some β-blockers can promote activation of β_2_AR and selectively transduce G protein/cAMP/PKA signaling but not GRK signaling. The β_2_AR-induced cAMP signal is highly restricted to the local domain, which selectively promotes activation of receptor-associated LTCC but not AMPAR in primary hippocampal neurons. Moreover, we have engineered a mutant β_2_AR that is selectively activated by β-blockers but not by catecholamines at low concentration. This study defines CAR and ALP as Gs-biased partial agonists of βAR for spatially highly restricted cAMP/PKA signaling to Ca_V_1.2 in neurons.

PKA-mediated phosphorylation is thought to play critical roles in heterologous desensitization of GPCRs and in receptor switching from Gs to Gi coupling (40, 41), whereas GRK-mediated phosphorylation is implicated in β-arrestin recruitment and β-arrestin-dependent ERK activation (13-18). We have recently characterized that PKA and GRKs phosphorylate distinct subpopulations of β_2_AR in a single fibroblast or neuron (12). While GRK phosphorylation of β_2_AR is only observed at high concentrations of agonists, PKA phosphorylation can be induced with minimal doses of agonist (12, 27, 28, 42). Here, our data show CAR does not promote GRK phosphorylation at low concentrations and induces a slow and minimal GRK effect at high concentrations when compared to those induced by ISO. The CAR-induced GRK effects are minimally related to the PKA effects. Previously, CAR has been recognized as a biased β-blocker that preferentially activates β-arrestin/ERK pathways (29, 43). Despite the prominent role of GRK phosphorylation in full agonist ISO-induced β_2_AR-β-arrestin/ERK signaling, our data clearly indicate that GRK phosphorylation of β_2_AR is not necessary for CAR-induced activation of ERK, consistent with a recent study showing a distinct general mechanism of β-arrestin activation that does not require the GRK-phosphorylated tail of different GPCRs (44). Meanwhile, other studies show that in the absence of all G proteins, GPCRs fail to transduce β-arrestin/ERK signaling (45). These data indicate the necessity of G proteins in GPCR-induced arrestin activation. In our study, we observed that a concentration-dependent correlation between PKA phosphorylation of β_2_AR with ERK activity induced by β-blockers, suggesting the potential role of Gs and PKA in CAR-induced β_2_AR-β-arrestin/ERK signaling are overlooked. In comparison, Gi is not required for CAR-induced β_2_AR/β-arrestin signaling even though CAR induces Gi recruitment to β_1_AR for transducing β_1_AR/β-arrestin signaling (46). Moreover, our results are also in line with a recent report that activation of β_2_AR with as low as femtomolar concentrations of ligands causes sustained ERK signaling (47), further support a PKA but GRK-dependent mechanism in GPCR-induced ERK activation. Future studies will help us understand how ligand-induced GPCRs utilize distinct mechanisms in activating β-arrestin/ERK pathway.

Engineered GPCRs have been widely applied in investigating structural and biological processes and behaviors by precisely controlling specific GPCR signaling branches (46). Previous mutagenesis studies have shown that β_2_AR with S204/207A mutation loses binding to adrenaline but still binds with several β-blockers including ALP (37). Based on this and recent advances in βAR structures with agonists and β-blockers (38, 39), we have generated a S204/207A mutant that bestow β_2_AR with the ability to be selectively activated by β-blockers such as CAR and to transduce cAMP/PKA signaling. At nanomolar concentrations, while ISO fails to stimulate PKA phosphorylation of the S204/207A mutant β_2_AR, the mutant receptor still retains CAR-induced stimulation of PKA-phosphorylation of the receptor. The CAR-induced activation of mutant β_2_AR triggers the β_2_AR/Gs/cAMP/PKA signaling pathway and selectively targets downstream effectors in primary hippocampal neurons. Interestingly, the S204/207A β2AR mutant is not only refractory to its agonists but also completely lost both ISO- and CAR-induced GRK-phosphorylation of β_2_AR. Further studies comparing this mutant with previous reported β_2_AR-TYY and Y219A mutants that lack Gs and GRKs coupling, respectively (18, 47), will facilitate the analysis of the physiological relevance of Gs/cAMP/PKA-dependent and GRK-dependent signaling pathways and enable researchers to explore β-arrestin/ERK pathway devoid of individual signaling branches.

β-blockers are a standard clinical treatment in a broad range of diseases. Many β-blockers possess intrinsic sympathomimetic activities (19, 20), which are problematic due to the side effects through stimulation of βARs (19, 20), a feature that limits the clinical utility of the drugs. Here, we show that β-blockers promote activation of β_2_AR by recruiting Gs that selectively transduces cAMP/PKA signal but not GRK signal. Meanwhile, binding of β-blockers to β_1_AR has been shown to enhance cAMP levels locally by dissociating a β_1_AR-PDE4 complex, thereby reducing the local cAMP-hydrolytic activity (48), β_1_AR and β_2_AR thus could utilize different mechanisms for β-blocker-induced signaling. Another interesting observation is that the β-blocker-induced β_2_AR-cAMP signal is sufficient to promote PKA phosphorylation of both β_2_AR and the receptor-associated Ca_V_1.2 of LTCC, but not another substrate, the AMPAR GluA1 subunit. Both LTCC and AMPAR are shown to associate with the β_2_AR in hippocampal neurons (30-33). Therefore, the preference of one local membrane target over another local target indicates a highly restricted nature of the cAMP-PKA activities, potentially dependent on the recently identified distinct subpopulations of β_2_AR and associated signaling molecules in the neurons (12). Nevertheless, the PKA phosphorylation leads to augmentation of LTCC activity, potentially contributing to the neuronal toxicities. Therefore, activation of GPCR at low ligand concentrations should be taken into consideration when designing and screening new therapeutic drugs.

## Methods

### Plasmids

DNA constructs expressing FLAG-tagged human β_2_AR (FLAG-β_2_AR) and HA-tagged rat L-type calcium channel (LTCC) α_1_1.2 were described before (12). FLAG-tagged human β_2_AR with S204/207A double mutations (FLAG-mutant) was generated by Gibson assembly method (Thermo Fisher) using FLAG-β_2_AR and synthetic gBlocks with the double mutations as templates (Integrated DNA Technologies). FRET biosensor ICUE3 was described before (34). To make the β_2_AR-ICUE3 fusion biosensor, ICUE3 was fused to the C-terminal of FLAG-β_2_AR with Gly-Ser linker. HA-Gsα was made by replacing CFP with HA tag, using Gsα-CFP as template (a gift from Dr. Catherine Berlot, Addgene plasmid # 55793).

### Antibodies and Chemicals

Mouse monoclonal antibodies against β_2_AR at serine 261/262 (clone 2G3) and at serine 355/356 (clone 10A5) were kindly provided by Dr. Richard Clark (UT Huston). Polyclonal antibodies against β_2_AR (sc-570) and phosphorylated β_2_AR at serine 355/356 (sc-16719R) were purchased from Santa Cruz Biotechnology. Polyclonal antibodies against α_1_1.2 residues 754-901 for total α_1_1.2 (FP1), residues 1923-1935 for phosphorylated serine 1928 site (LGRRApSFHLECLK, pS1928) and residues 1694-1709 for phosphorylated serine 1700 site (EIRRAIpSGDLTAEEEL, pS1700) were described before (49). Polyclonal antibodies against GluA1 residues 894-907 for total GluA1, residues 826-837 for phosphorylated serine 831 site (LIPQQpSINEAIK, pS831) and residues 840-851 for phosphorylated serine 845 site (TLPRNpSGAGASK, pS845) were from Cell Signaling (Danvers, MA). Other antibodies used in the experiments include: anti-FLAG (F3040, Sigma), anti-HA (MMS-101R, Covance), Alexa fluor 488 conjugated goat anti-rabbit IgG and Alexa fluor 594 conjugated goat anti-mouse IgG (A-11034 and A-11032, Thermo Fisher), DyLight 680 conjugated goat anti-mouse IgG and anti-rabbit IgG (35518 and 35568, Thermo Fisher), IRDye 800CW conjugated goat anti-mouse IgG and anti-rabbit IgG (926-32210 and 926-32211, Li-cor).

Isoproterenol (I2760), timolol (T6394), alprenolol (A8676), propranolol (P0884), metoprolol (M5391), CGP12177A (C125), CGP20712A (C231), ICI118551 (I127), 3-isobutyl-1-methylxanthine (I5879) and 2’,5’-dideoxyadenosine (D7408) were purchased from Sigma. Carvedilol (15418) was from Cayman Chemical, H89 (H-5239) was from LC Labs, pertussis toxin (179B) was from List Labs.

### Cell Culture and Transfection

Human embryonic kidney HEK293 cells were from American Type Culture Collection (ATCC) and were maintained in Dulbecco’s modified Eagle medium (DMEM, Corning) supplemented with 10% fetal bovine serum (FBS, Sigma). HEK293 cells stably expressing FLAG-β_2_AR was from previous study (35). HEK293 cells stably expressing FLAG-mutant β_2_AR was generated in this study. Briefly, cells transfected with β_2_AR-mutant were selected by G418 resistance (Corning) and cell clones were obtained by limiting serial dilution, monoclonal cells were analyzed by western blots and the one with comparable β_2_AR expression to FLAG-β_2_AR stable cells was chosen. Mouse embryonic fibroblasts (MEFs) from β_1_AR/β_2_AR double knockout (DKO) mouse was described in previous study (50) and were maintained in DMEM supplemented with 10% FBS. Primary mouse hippocampal neurons were isolated and cultured from P0-P1 early postnatal DKO mouse pups, and primary rat hippocampal neurons were prepared from E17-E19 embryonic rats using methods described previously (51, 52). Briefly, dissected hippocampi were dissociated by 0.25% trypsin (Corning) and trituration. Neurons were plated on poly-D-lysine-coated (Sigma) glass coverslips in 24-well plate for imaging and in 6-well plate for biochemistry at a cell density of 50,000/well and 1 million/well, respectively. Neurons were cultured in Neurobasal medium supplemented with GlutaMax and B-27 (Thermo Fisher). Animal protocols were approved by IACUC of the University of California at Davis according to NIH regulations.

HEK293 cells were transfected with plasmids using polyethylenimine according to manufacturer’s instructions (Sigma). Neurons were transfected by the Ca^2+^-phosphate method (53). Briefly, cultured neurons on 6-10 DIV were switched to pre-warmed Eagle’s minimum essential medium (EMEM, Thermo Fisher) supplemented with GlutaMax 1 hour before transfection, conditioned media were saved. DNA precipitates were prepared by 2x HBS (pH 6.96) and 2 M CaCl_2_. After incubation with DNA precipitates for 1 hour, neurons were incubated in 10% CO_2_ pre-equilibrium EMEM for 20 minutes, then replaced with conditioned medium and cultured in 5% CO_2_ incubator until use.

### Confocal Microscopy Imaging

Rat hippocampal neurons were transfected with FLAG-β_2_AR on 10 DIV, treated for 5 minutes with 10 nM or 1 μM indicated drugs on 12 DIV. Mouse DKO hippocampal neurons were transfected with FLAG-β_2_AR or FLAG-mutant and HA-α_1_1.2 at 1:1 ratio on 6-8 DIV, and stimulated with indicated drugs and times 24 hours after transfection. Treated cells were fixed, permeabilized, and co-stained with indicated antibodies with a final concentration of 1 μg/ml for each antibody, which were revealed by a 1:1000 dilution of Alexa fluor 488 conjugated goat anti-rabbit IgG or Alexa fluor 594 conjugated goat anti-mouse IgG, respectively. Fluorescence images were taken by Zeiss LSM 700 confocal microscope with a 63×/1.4 numerical aperture oil-immersion lens.

### Proximity ligation assay

HEK293 cells growing on poly-D-lysine coated coverslips were transfected with FLAG-β_2_AR or FLAG-mutant, HA-Gsα and pEYFP-N1 at 8:1:1 ratio. 24 hours after transfection, cells were serum-starved 2 hours, treated 100 nM indicated drugs for 5 minutes. Following stimulation, cells were fixed, permeabilized, and co-stained with anti-β_2_AR antibody (1:100 dilution) from rabbit in conjunction with anti-HA antibody (1:1000 dilution) from mouse. The proximity ligation reaction was performed according to the manufacturer’s protocol using the Duolink in situ detection orange reagents (Sigma). Images were recorded with Zeiss LSM 700 confocal microscope with a 63×/1.4 numerical aperture oil-immersion lens. To quantify the PLA signals, the number of red fluorescent objects in each image was quantified using the Squassh plug-in for ImageJ software (54), and divided by the number of transfected cells.

### Fluorescence resonance energy transfer (FRET) measurement

FRET measurement was performed as previously described (35). Briefly, HEK 293 cells were transfected with ICUE3 or β_2_AR-ICUE3, DKO MEFs were co-transfected with ICUE3 and FLAG-β_2_AR or FLAG-mutant. Cells were imaged on a Zeiss Axiovert 200M microscope with a 40×/1.3 numerical aperture oil-immersion lens and a cooled CCD camera. Dual emission ratio imaging was acquired with a 420DF20 excitation filter, a 450DRLP diachronic mirror, and two emission filters (475DF40 for cyan and 535DF25 for yellow). The acquisition was set with 0.2 second exposure in both channels and 20 second elapses. Images in both channels were subjected to background subtraction, and ratios of yellow-to-cyan were calculated at different time points.

### Western blot

HEK293 cells stably expressing FLAG-β_2_AR or FLAG-mutant were serum-starved for 2 hours and treated with indicated drugs and times, then harvested by lysis buffer (10 mM Tris pH 7.4, 1% NP40, 150 mM NaCl, 2 mM EDTA) with protease and phosphatase inhibitor cocktail. Rat hippocampal neurons on 10-14 DIV were treated with indicated drugs and times, then harvested by lysis buffer (10 mM Tris pH 7.4, 1% TX-100, 150 mM NaCl, 5 mM EGTA, 10 mM EDTA, 10% glycerol) with protease and phosphatase inhibitor cocktail. Protein samples were analyzed by Western blot using antibodies as indicated at a 1:1000 dilution and signals were detected by Odyssey scanner (Li-cor). Uncropped scans are presented in Supplemental Fig. 6.

### Cell-attached Patch Clamp Electrophysiology

Primary rat and mouse hippocampal neurons were used on 7-10 DIV. Cell-attached patch clamp recordings were performed on an Olympus IX70 inverted microscope in a 15-mm culture coverslip at room temperature (22-25 °C). Signals were recorded at 10 kHz and low-pass filtered at 2 kHz with an Axopatch 200B amplifier and digitized with a Digidata 1440 (Molecular Devices). Recording pipettes were pulled from borosilicate capillary glass (0.86 OD) with a Flaming micropipette puller (Model P-97, Sutter Instruments) and polished (polisher from World Precision Instruments). Pipette resistances were strictly maintained between 6-7 MΩ to ameliorate variations in number of channels in the patch pipette. The patch transmembrane potential was zeroed by perfusing cells with a high K^+^ extracellular solution containing (in mM) 145 KCl, 10 NaCl, and 10 HEPES, pH 7.4 (NaOH). The pipette solution contained (in mM) 20 tetraethylammonium chloride (TEA-Cl), 110 BaCl_2_ (as charge carrier), and 10 HEPES, pH 7.3 (TEA-OH). This pipette solution was supplemented with 1 µM ω-conotoxin GVIA and 1 µM ω-conotoxin MCVIIC to block N and P/Q-type Ca^2+^ channels, respectively, and (S)-(-)-BayK-8644 (500 nM) was included in the pipette solution to promote longer open times and resolve channel openings. Indeed, BayK-8644 is routinely used to augment detection of L-type channels in single-channel recordings (49, 55, 56). To examine the effects of β-adrenergic stimulation on the L-type Ca_V_1.2 single-channel activity, 1 μM isoproterenol was added to the pipette solution in independent experiments. Note that we have previously used the L-type Ca_V_1.2 channel blocker nifedipine (1 μM) to confirm the recording of L-type Ca_V_1.2 currents under control conditions and in the presence of isoproterenol (49). Single-channel activity was recorded during a single pulse protocol (2 seconds) from a holding potential of −80 mV to 0 mV every 5 seconds. An average of 50 sweeps were collected with each recording file under all experimental conditions. The half-amplitude event-detection algorithm of pClamp 10 was used to measure overall single-channel L-type Ca_V_1.2 activity as nPo, where n is the number of channels in the patch and Po is the open probability. Note that the number of channels in each patch recording (n) was not estimated and that all data are presented as “nPo” (product of n and channel open probability). nPo values were pooled for each condition and analyzed with GraphPad Prism software.

## Statistical analysis

Data were analyzed using GraphPad Prism software and expressed as mean ± s.e.m. Differences between two groups were assessed by appropriate two-tailed unpaired Student’s t-test or nonparametric Mann-Whitney test. Differences among three or more groups were assessed by One-way ANOVA with Tukey’s post hoc test. P < 0.05 was considered statistically significant.

## Data availability

The data that support the findings of this study are available from the corresponding author upon reasonable request.

## Acknowledgements

This work was supported by NIH grant GM129376 and VA Merit grant BX002900 to Y.K.X., HL098200 and HL121059 to M.F.N. A.S. and Q.S. were recipients of AHA postdoctoral fellowship. Y.K.X. is an established AHA investigator.

## Author contributions

A.S. and Y.K.X. conceived and designed experiments. A.S. generated DNA constructs and stable cells, did mouse neuron culture, imaging, FRET and Western blot. M.F.N. performed single channel recording. D.C., B.X. and J.M.M helped Western blot. M.K and Q.S. helped DNA constructs and FRET. K.M.M provided rat neuron culture. A.S. and Y.K.X. interpreted all the data and wrote the manuscript with inputs from X.-Y.Y., J.W.H. and M.F.N. Y.K.X. provided overall project supervision.

